# The Surgical Method of Craniectomy Differentially Affects Acute Seizures, Brain Deformation and Behavior in a TBI Animal Model

**DOI:** 10.1101/2024.06.28.601257

**Authors:** Cesar Santana-Gomez, Gregory Smith, Ava Mousavi, Mohamad Shamas, Neil G. Harris, Richard Staba

**Author notes:** **Corresponding Author:** Richard Staba, Department of Neurology, 710 Westwood Plaza, RNRC 2145, Los Angeles, CA, US 90095. Department of Neurology, 710 Westwood Plaza, RNRC 2144-1, Los Angeles, CA, US 90095, Department of Neurosurgery, 300 Stein Plaza, STE 551, Los Angeles, CA 90095, Department of Neurology, 710 Westwood Plaza, RNRC 2144, Los Angeles, CA, US 90095, Department of Neurology, 710 Westwood Plaza, RNRC 2144-2, Los Angeles, CA, US 90095, Department of Neurosurgery, 300 Stein Plaza, STE 551, Los Angeles, CA 90095.

## Abstract

Traumatic brain injury (TBI) is the leading cause of morbidity and mortality worldwide. Multiple injury models have been developed to study this neurological disorder. One such model is the lateral fluid-percussion injury (LFPI) rodent model. The LFPI model can be generated with different surgical procedures that could affect the injury and be reflected in neurobehavioral dysfunction and acute EEG changes. A craniectomy was performed either with a trephine hand drill or with a trephine electric drill that was centered over the left hemisphere of adult, male Sprague Dawley rats. Sham craniectomy groups were assessed by hand-drilled (ShamHMRI) and electric-drilled (ShamEMRI) to evaluate by MRI. Then, TBI was induced in separate groups (TBIH) and (TBIE) using a fluid-percussion device. Sham-injured rats (ShamH/ShamE) underwent the same surgical procedures as the TBI rats. During the same surgery session, rats were implanted with screw and microwire electrodes positioned in the neocortex and hippocampus and the EEG activity was recorded 24 hours for the first 7 days after TBI for assessing the acute EEG seizure and Gamma Event Coupling (GEC). The electric drilling craniectomy induced greater tissue damage and sensorimotor deficits compared to the hand drill. Analysis of the EEG revealed acute seizures in at least one animal from each group after the procedure. Both TBI and Sham rats from the electric drill groups had a significant greater total number of seizures than the animals that were craniectomized manually (p<0.05). Similarly, EEG functional connectivity was lower in ShamE compared to ShamH rats. These results suggest that electrical versus hand drilling craniectomies produce cortical injury in addition to the LFPI which increases the likelihood for acute post-traumatic seizures. Differences in the surgical approach could be one reason for the variability in the injury that makes it difficult to replicate results between preclinical TBI studies.

## Introduction

The lateral fluid percussion injury (LFPI) is a well-studied model of human traumatic brain injury (TBI)^1–3^, and has been used in rabbits, cats, rats, mice, and pigs ^4–8^. The injury is produced by applying a short duration fluid pulse (∼20 msec) through a craniectomy against an intact, exposed dura, resulting in a transient deformation of the brain. The LFPI model can produce graded neurological, histological, and cognitive outcomes that replicate those seen in human TBI^9,10^, and in cases of moderate to severe LFPI, acute and sometimes late posttraumatic seizures^11,12^.

As with all experimental models, modifications are made to the LFPI rat model depending on the surgeon’s technique and experience, and/or goals of the research. For example, some studies report performing the craniectomy with a dental electric drill adapted with jeweler’s bur or trephine^12,13^, while others report using a hand drill trephine^14^. Studies show an extensive network of blood vessels and nerves between the rat brain and calvaria^15^ and craniectomy alone, especially when performed with an electrical drill, can disrupt this network, resulting in an upregulation of pro-inflammatory proteins^16–18^, leaky blood vessels^18–20^, and cerebral edema^17^. Under these conditions it is reasonable to predict LFPI with an electric-drilled craniectomy, which can generate measurable heat, vibration, and axial force to the underlying tissue^18^, would be more severe than one with a hand-drilled craniectomy. However, the potential differences in severity during the acute period after LFPI with respect to Magnetic Resonance Imaging (MRI) morphology, electroencephalograph (EEG) functional changes, and behavior is not well-documented.

To address this knowledge gap in the LFPI model, in the current study we performed a series of experiments to first evaluate the MRI morphological and EEG functional connectivity changes associated with an electric-drilled versus hand-drilled craniectomy. Then, in separate cohorts of rats, we performed the same craniectomy procedures in LFPI and sham-injured rats and assessed each for sensorimotor performance up to 28 days after injury and acute posttraumatic seizures during the first 7 days after injury.

## Material and Methods

### Experimental Overview

Adult male Sprague-Dawley rats (n = 50, 300–350g at the time of the craniectomy; Charles River Laboratories Inc.) were randomized into two groups for MRI only and 4 groups for EEG studies (Fig. 1). The rats were housed in individual cages in a controlled environment (temperature, 21 – 26 °C, humidity 30-70 %, lights on 06:00–18:00 h) and had free access to food and water. All animal procedures were approved by the University of California Los Angeles Institutional Animal Care and Use Committee (protocol 2000- 153- 61 A).

**Fig.1.**
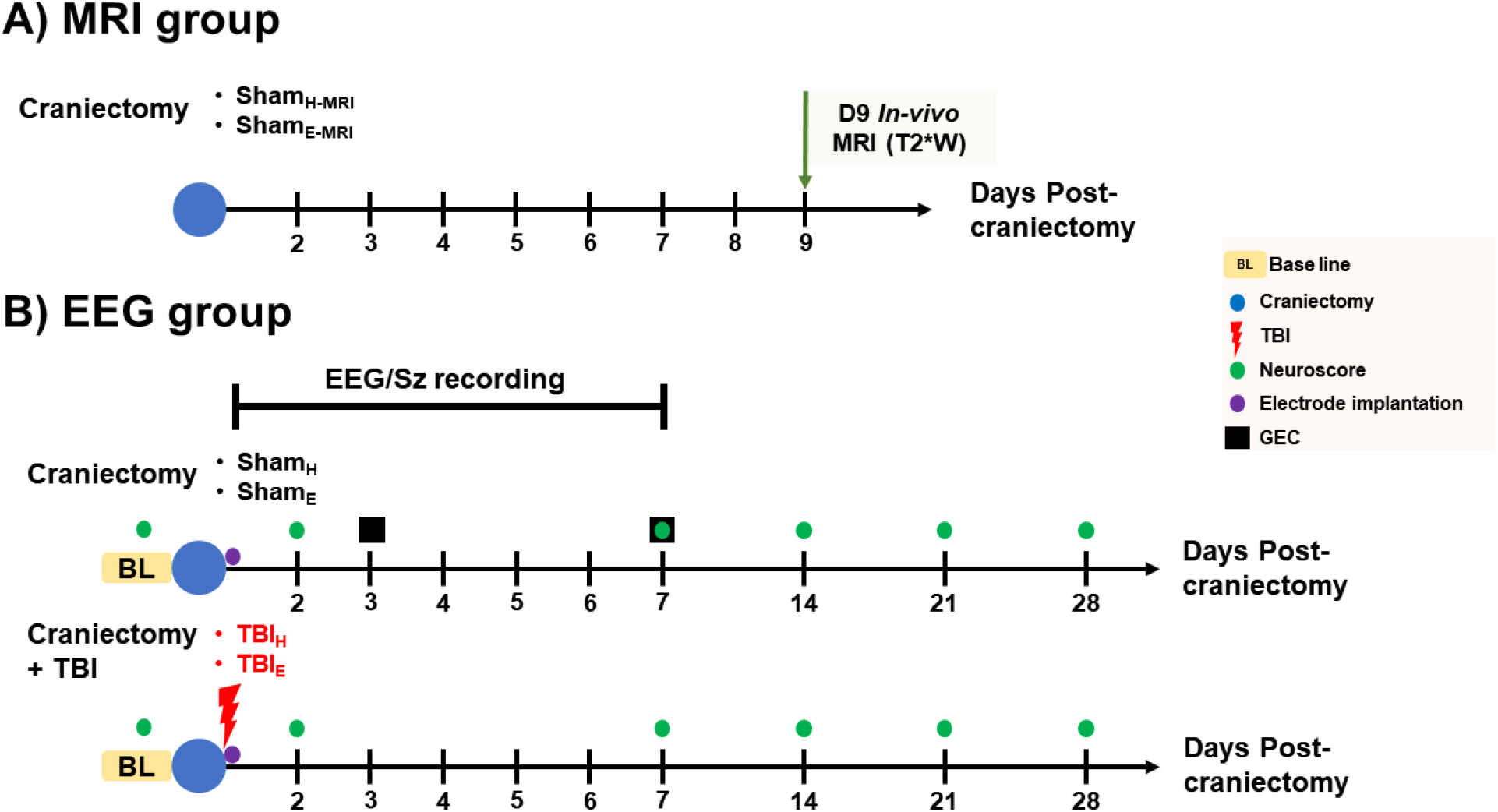
Experimental groups and timeline. The rats were randomized into MRI and EEG groups. A) The MRI group assessed the brain structural changes induced by two different craniectomy methods (denoted by blue circle on timeline) on sham operated animals (ShamH-MRI, ShamE-MRI). The animals were imaged on day 9 post-craniectomy. B) In the EEG group, the effect of craniectomy on EEG function connectivity was assessed (top part; ShamH, ShamE). Electrodes (purple circle) were implanted after craniectomy during same surgery. Functional connectivity was computed as gamma event connectivity (GEC; black square) on days 3 and 7 post-craniectomy. In a separate set of experiments, the effects of craniectomy and TBI on sensorimotor deficits (Neuroscore; green circles) and early seizures (bottom part; TBIH, TBIE). Neuroscore tests were performed at 6 time points, including baseline (BL) and EEG was reviewed for early seizures during first 7 days post-craniectomy.

### Surgery

All rats were anesthetized using 5% isoflurane. The fur atop the rat’s head was shaved, the skin scrubbed with antiseptic, and the rat was draped for surgery. A 3-4 cm midline incision was made, with the skin parted, and the exposed underlying skull was cleaned and dried. In all cases, the craniectomy was positioned over the left parietal bone and centered AP −4.5 mm from bregma and ML +2.5 mm from the sagittal suture (Fig. 2)^21^.The craniectomy was performed using a stainless-steel trephine drill bit (Ø 5 mm; Meisinger, Germany) that was either mounted to a fine pin vise hand drill (Tamiya, Inc., Japan) or single-speed Dremel MultiPro electric drill (35,000 r.p.m; Dremel US, Mt. Prospect, IL). Trephination used light, but steady, pressure and alternated between drilling and irrigating with room temperature sterile saline to remove bone dust and reduce bone heating, and was completed in about 30 sec. After the bone was removed, the dura was inspected to ensure it was intact, and the incision was closed using non-absorbable surgical suture (4-0 nylon suture, Ethicon). The surgical procedures for the MRI and EEG cohorts were performed by two surgeons (CSG and GS) with comparable experience who had gone through the same surgical trainings, ensuring consistency in the procedures performed.

**Fig. 2.**
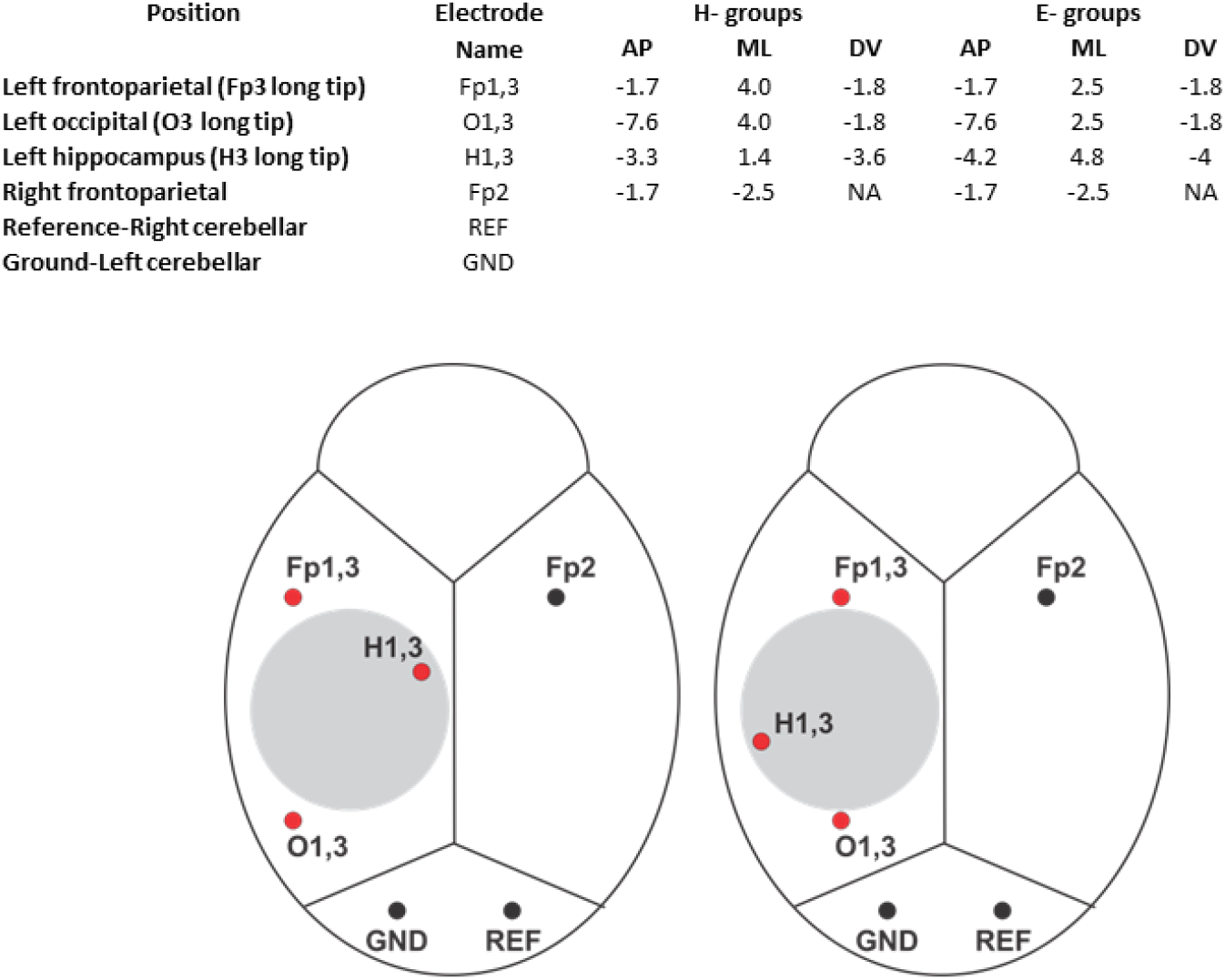
Electrode placements for EEG recording (adapted from Paxinos and Watson, 2007)^21^. Coordinates associated with each craniectomy method, which were slightly modified according to the experimental protocol (upper panel). Electrodes were positioned anterior and posterior to the Ø 5 mm craniectomy (grey circle, lower panel), within hippocampus, and over the contralateral hemisphere. Abbreviations: Fp1,3, bipolar ipsilateral frontoparietal; O1,3, bipolar ipsilateral occipital; H1,3, bipolar hippocampus; Fp2, Contralateral frontoparietal; red dots, bipolar penetrating electrodes; black dots, screw electrodes; GND, Ground; REF; Reference.

### MRI acquisition, post-processing, and analysis

To assess the effects of the craniectomy on brain tissue, MRI studies were performed 9 days after surgery in 5 rats that had craniectomy with hand drill (ShamH-MRI) and another 5 rats with craniectomy using the electrical drill (ShamE-MRI). MRI images were acquired using a Bruker Biospin, 7T scanner and a 400 mT/m gradient with a 110 µs rise-time and an actively decoupled, quadrature, surface coil. For anatomical scans a multi-echo, gradient echo (MGE), 3-dimensional sequence was used with a TR of 125 ms, 13 echos and an effective TE of 2.8 ms to 52 ms with a 4.08 ms spacing, a 20° flip angle, and a data matrix of 160 X 122 X 80 mm. Resulting image resolution was 160 mm^3^ isometric. The 13 echoes were averaged to create a T2* weighted map for each rat which was normalized for signal intensity and used to generate a population mean deformation template from rats in all groups using ANTs^22^. The warp transformation parameters generated by this procedure were then used to obtain a Jacobian determinant map for each rat that reflects local tissue deformation because of the sham or injury procedures. Regions of greatest increase or decrease in tissue formation were found in each brain using population-based, voxel-based statistics by defining a boundary Z score threshold at Z>2.33 (p<0.01) calculated using the average ShamH-MRI Jacobian determinant maps and its standard deviation. The ShamH-MRI group was chosen because our hypothesis and data suggested this group would have the least tissue deformation. The voxels surviving this threshold were binarized and summed across all rats/group to create the population count overlay map (Fig. 3D). The volume of thresholded voxels was taken from each of the ShamE-MRI rats as a measure of significant positive tissue deformation when compared to the amount of tissue deformation in the ShamH-MRI rats.

**Fig. 3.**
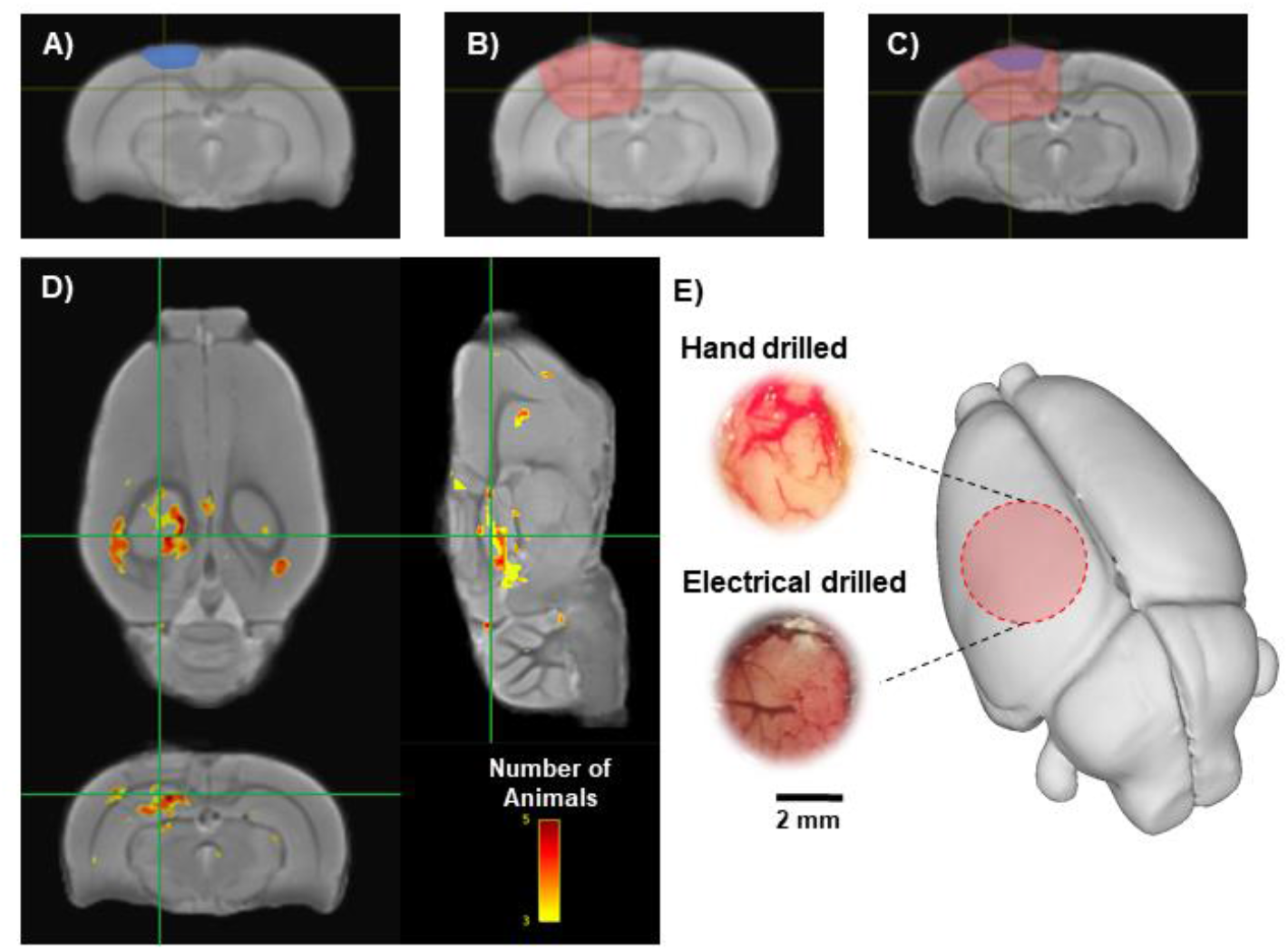
T2* weighted MRI comparison in sham-injured rats. A-B) The depth of tissue damage highlighted over the mean T2* weighted images of the hand- (A; blue-shaded area) and (B; red-shaded area) electrical-drilled craniectomy. C) Overlap of tissue damage between the hand- and electrical-drilled craniectomy superimposed on the mean template of all brains. D) Images illustrate brain regions associated with positive tissue local tissue deformation that was greater in electrical-drilled than hand-drilled craniectomies. Color intensity represents the number of brains altered in that region. Background image is a T2* weighted mean of all brains used in this comparison (ShamHMRI, n=5; ShamEMRI, n=5). E) Macroscopic view after craniectomy method by hand (top) or electrical drill (bottom) craniectomy method in sham-injury animals. Photographs show darker dura and blood vessels within the craniectomy produced by electrical drill compared to hand drill. Red circle represents the position of craniectomy site.

### Induction of lateral fluid percussion injury (LFPI)

To evaluate the effects of craniectomy in the LFPI model of human TBI, craniectomies were performed (see surgery section) in 13 rats that had craniectomy with hand drill (TBIH) and another 14 rats with craniectomy using the electrical drill (TBIE). For the LFPI induction a modified Luer-Lock syringe cap was anchored over the craniectomy and set with dental acrylate. A severe TBI was induced with a fluid percussion injury pendulum device equipped with a straight-tip attachment (AmScien Instruments, Model FP 302, Richmond, VA, USA,)^23^. Sham-injured control rats for hand drill (ShamH, n=6) and electrical drill (ShamE, n=7) craniectomies underwent the same surgical procedures as the TBI rats, but LFPI was not performed.

### Microelectrode Implantation and EEG recording

EEG recordings were performed in the TBI and Sham groups using microelectrodes and epidural screw electrodes implanted during the same surgery for LFPI. Paired microwires (Ø 40 µm) with tip separation of 0.5 mm were implanted in the perilesional fronto-parietal cortex (Fp1,3), posterior to the craniectomy in occipital cortex (O1,3), and ipsilateral anterior hippocampus (H1,3). One screw electrode was implanted into contralateral fronto-parietal cortex (Fp2). Ground (GND) and reference (REF) electrodes (stainless steel screws) were inserted in the skull bone overlying the cerebellum. Electrode locations are summarized in Fig. 2. The electrodes were mounted in a 12-channel Plastics One pedestal (M12P, PlasticsOne Inc.). The pedestal was secured to the skull with dental acrylic. Sham-injured controls underwent the same electrode implantation procedures as TBI rats.

### EEG Data Acquisition

Immediately following surgery, the rats were placed in a Plexiglas cage with temperature-regulated (∼37 °C) and watched during recovery before being returned to the home Plexiglas cylinder (about 30 min). The electrode headset atop the rat skull was connected to a 12-pin swivel commutator (SL12C; PlasticsOne Inc.) via a flexible shielded cable (M12C-363/2; PlasticsOne Inc.), allowing the rat to move freely during recording. The commutator was connected to an amplifier using a flexible shielded cable 363/2- 441/12 (PlasticsOne Inc.). Electrical brain activity was recorded using an Intan RHD200 amplifier and began within an hour after completion of the surgery and continued 24 h/day for the first week after TBI. EEG was recorded with respect to REF electrode overlying the cerebellum and sampled at a minimum of 2 kHz per channel and band-passed between 0.1 Hz and 1 kHz.

### Post-injury functional impairment evaluation

Post-injury sensorimotor deficits were assessed based on the composite neuroscore as previously described^24^. Rat responses were nominally scored from 0 (severely impaired) to 4 (normal) for (i) left and right contraflexion, (ii) left and right hindlimb flexion, (iii) left and right lateral pulsion, and (iv) ability to stand on an inclined board in a vertical and horizontal (left and right) position. The maximum possible score was 28. Baseline neuroscore was performed 2 days prior to injury and then 2 days after injury and then each week for the first month. For the baseline inclined board assessment, the angle at which the rat was able to keep a steady posture was given a maximal score of 4. For the post-injury incline board assessment, the injury score was computed as the difference in angle between post-injury and baseline periods (4 = no difference in the angle; 3=2.5° less than the baseline; 2=5° less than the baseline; 1=7.5° less than the baseline; 0=10° less than the baseline) ^24^.

### Seizure detection

The EEG recordings were exported into European data format (.EDF) and then analyzed using EDF browser software (https://www.teuniz.net/edfbrowser/). EEG was reviewed using a 90 Hz low pass filtered displayed in 2 minutes windows. An acute (<=7 days post-injury) electrographic seizure was defined as an event consisting of repetitive epileptiform EEG spike discharges >2 Hz with quasi-rhythmic, spatial, and temporal changes in frequency, amplitude, morphology lasting at least 10 seconds^25^.

### Data analysis for Gamma Event Coupling

Gamma event coupling (GEC) in ShamH and ShamE was computed from a 10- minute episode of EEG with high-amplitude, irregular activity typical of slow wave sleep on days 3 and 7 after surgery. Episodes were selected that were free of movement or obvious muscle-related artifacts or channels with power line noise. Briefly, the EEG was down sampled to 1 kHz and bandpass filtered between 30 and 55 Hz. Then, the maximum peak of each gamma wave was detected and the lead or lag between gamma maxima from each unique pair of electrodes was quantified using peri-event histograms. Shannon entropy was used to evaluate each histogram and a connectivity factor was calculated by subtracting the observed entropy from the computed maximum entropy and dividing the difference by the computed maximum entropy. These calculations produced a connectivity factor between 0 and 1, where 0 means fully disconnected and 1 means fully connected. Connectivity factors were arranged in symmetrical connectivity matrices where the intersection between a row and column represent the connectivity between a pair of electrodes. To compensate for data loss of noisy channels, a recovery algorithm was used to interpolate the connectivity factor if (1) only one electrode was missing in a rat and day, (2) this electrode was missing in day 3 or 7 but not both days, and (3) data from the electrode was available for at least 2 rats in the same group on both days. To recover the missing data, the percentage of change in connectivity strength between day 3 and 7 was calculated for the missing electrode in all rats in the same group. Since the data was available for one day in the considered rat, the average percentage of change was applied to interpolate the pairwise connectivity on the day when the electrode data was missing.

### Statistical analysis

Data analysis was performed using GraphPad Prism (V. 7.03). First, all data was assessed for normal distribution using D’Agostino & Pearson’s omnibus normality test. If not normally distributed, then a non-parametric test was used to compare the variable. The chi-square test was used to assess differences in post impact parameters and percentage of rats with acute EEG seizures. The differences in angle, pressure, apnea, and righting reflex were calculate using t-test. Neuroscore and seizure-related differences were analyzed using the one-way analysis of variance (ANOVA) test followed by post hoc analysis with Bonferroni correction. The GEC connectivity value of each pair of channels represented in the upper half of the connectivity matrix for rats in the ShamE group was compared to the corresponding pair of channels in the ShamH group using non-parametric Wilcoxon 2-tail test as the data was not normally distributed. The elements in the connectivity matrices were not treated as independent characteristics as they are coming from the same brain for each sample and hence the statistical comparison was corrected for multiple comparisons using FDR correction scheme to eliminate the possibility of statistical difference due to chance and to ensure that it’s directly related to the experimental protocol. The difference was considered significant at p ≤ 0.05.

## Results

### Craniectomy

Visual inspection of the brain’s surface through the craniectomy verified that the dura was intact in all rats (Fig. 3). In rats that had a craniectomy performed using a hand drill (ShamH and TBIH groups), the underlying brain tissue appeared light pink in color and the blood vessels presented a red color (Fig. 3E). In contrast, in rats that had a craniectomy using the electric drill (ShamE and TBIE groups), the underlying brain tissue had a dark red, brown hue and the blood vessels were dark red (Fig. 3E). There was no evidence of bleeding from large vessels on the brain surface in hand- or electric-drilled rats.

### MRI abnormalities after craniectomy

MRI was performed to evaluate the effects of hand- and electrical-drilled craniectomies on the underlying brain. The amount of positive tissue deformation was greater in the electrical-drilled craniectomy animals (ShamE-MRI) compared to the ShamH- MRI group (Figs. 3A-C). When compared to the ShamH-MRI group the average volume of tissue that was significantly deformed in the ShamE-MRI group was 34.7 ± 21.6 mm^3^. Deformed tissue was identified on a voxel-by-voxel basis and was considered deformed if the voxel had a z > 2.33 when compared to the ShamH-MRI group. The location of the significant voxels showing deformation was in an area directly under the craniectomy in most of the ShamE-MRI rats (Fig. 3D).

### Reduced EEG functional connectivity after electric-drilled craniectomy

In a separate cohort of sham-injured rats, gamma event connectivity (GEC; 30-55 Hz) was computed to assess the effects of the craniectomy procedure on EEG functional connectivity. Three days after surgery ShamE rats had weaker GEC between ipsilateral hippocampus and prefrontal cortex (H3-Fp1, p=0.03; H3-Fp3, p=0.01), as well as to occipital cortex (H3-O1, p=0.007; H3-O3, p=0.03), than ShamH rats (Fig. 4, uncorrected for multiple comparisons). Also, ShamE rats had weaker GEC between contralateral prefrontal cortex and ipsilateral occipital cortex than ShamH rats (Fp2-O3, p=0.001). Seven days after injury ShamE rats still had weaker GEC between ipsilateral hippocampus and prefrontal cortex (H3-Fp1, p=0.01), and between contralateral prefrontal cortex and ipsilateral occipital cortex (Fp2-O3, p=0.01) and prefrontal cortex (Fp2-Fp3 p=0.02) than the ShamH rats. At this same time ShamE rats had stronger GEC between ipsilateral hippocampus and contralateral prefrontal cortex than the ShamH rats (H3-Fp2, p=0.01; Fig. 4).

**Fig. 4.**
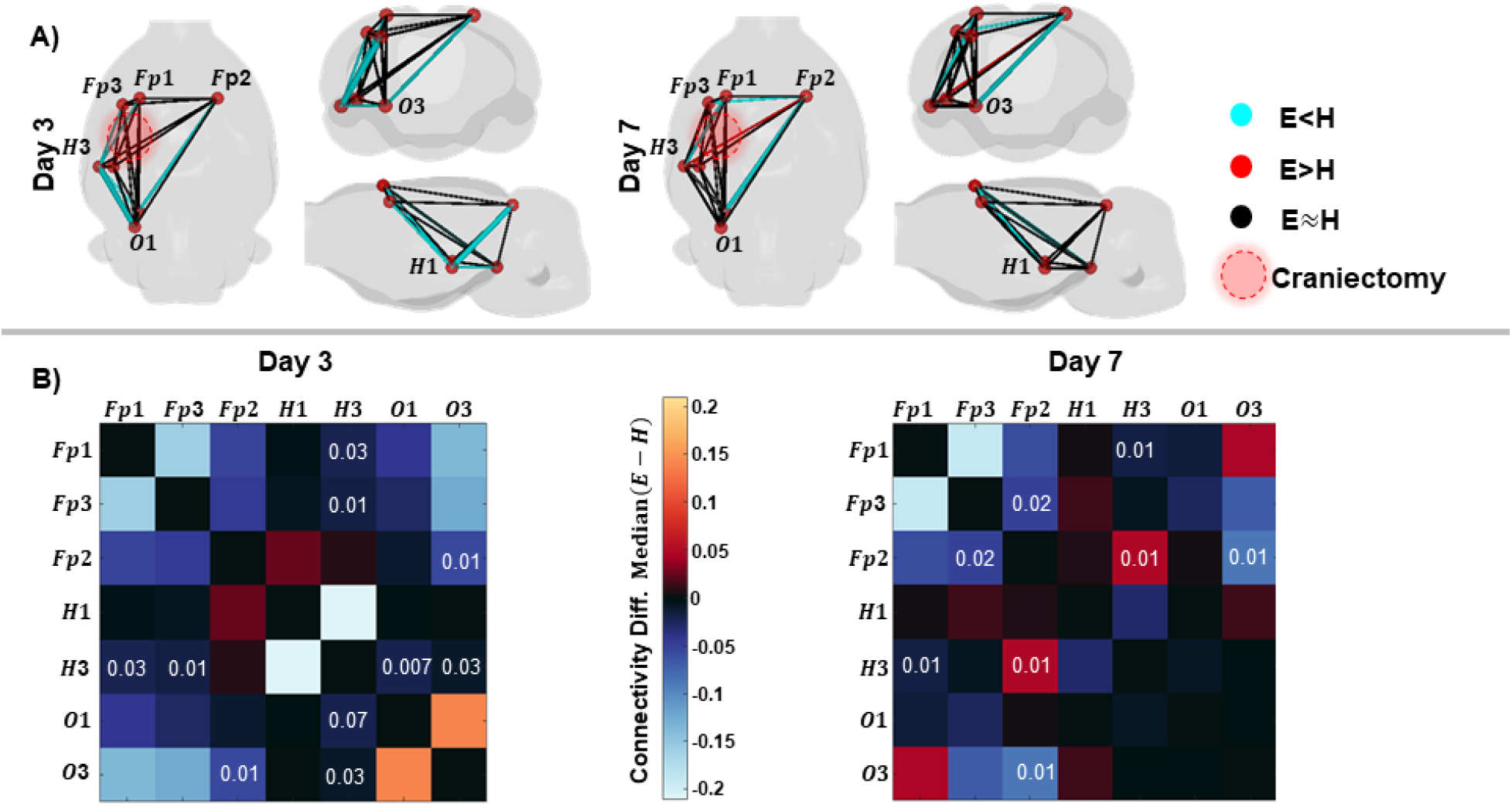
Differences in functional connectivity between ShamE and vs ShamH rats. A) Connectivity computed as GEC (30-55 Hz) between recording electrodes or nodes (red spheres) on days 3 (left) and 7 (right) after surgery. Differences in the median connectivity between ShamH and vs ShamE rats represented by the color and proportional to the thickness of lines between nodes. The cyan color denotes weaker connectivity in ShamE than ShamH group, while red denotes stronger GEC in ShamE than ShamH group, and black lines indicate no difference. Results depicted in axial, coronal and sagittal views. The injury site is represented as a translucent red circle on the axial view. B) Adjacency matrices of the differences in median connectivity between ShamE and ShamH groups on day 3 (left) and day 7 (right) after surgery. Value at the intersection of the row and column represent the p-value (less than 0.05) from the Wilcoxon test for corresponding electrode pairs. Note differences in connectivity were not statistically significant after FDR correction.

### Induction of Lateral Fluid Percussion Injury

The mean pendulum angle used for the LFPI to produce severe TBI was 19.8 ± 0.8° in TBIH rats and 17.8 ± 1.3° in TBIE rats (p<0.05; Table 1). The larger angle did not affect the impact pressure between the two groups (p>0.05) but was associated with a longer mean post-impact apnea (p<0.05). After recovery from impact, electrodes were implanted at the coordinates listed in Fig. 2. In sham-injured or TBI groups, there was no difference in mortality between hand- versus electric-drilled craniectomy. TBIH and TBIE groups showed higher proportion of rats that died after injury compared with ShamH animals (54% and 57% respectively, chi-square test p<0.05; Table 1).

**Table 1.** TBI rat model details.

| **Group** | **Angle**<br>**(°)** | **Pressure**<br>**(atm)** | **Apnea**<br>**(s)** | **Mortality rate**<br>**(%)** |
| --- | --- | --- | --- | --- |
| **Sham<sub>H</sub>** | --- | --- | --- | 0/6 (0%) |
| **Sham<sub>E</sub>** | --- | --- | --- | 1/7 (14.3%) |
| **TBI<sub>H</sub>** | 19.8±0.8 | 2.4±0.1 | 31.3±18.4 | 7/13 (53.9%)* |
| **TBI<sub>E</sub>** | 17.8±1.3<sup>&</sup> | 2.3±0.1 | 24.6±10.4<sup>&</sup> | 8/14 (57.2%)* |

\*p≤0.05 vs Sham<sub>H</sub>, Chi-square test; <sup>&</sup>p≤0.05 vs TBI<sub>H</sub>, t-test.

### Neuroscore Test

Assessment of sensorimotor function in craniectomized only rats revealed that there were relatively stable neuroscore values in ShamH rats throughout the duration of monitoring (p>0.05, Fig. 5). As expected, neuroscore values in TBIH rats significantly decreased 2 days post-injury (p<0.001; Fig. 5), started to recover toward baseline values at 7 days, and thereafter were like ShamH rats. By contrast, in ShamE and TBIE rats, neuroscore values were significantly lower than ShamH-injured rats at all time points, especially during the first 14 days after TBI (p<0.05; Fig. 5).

**Fig. 5.**
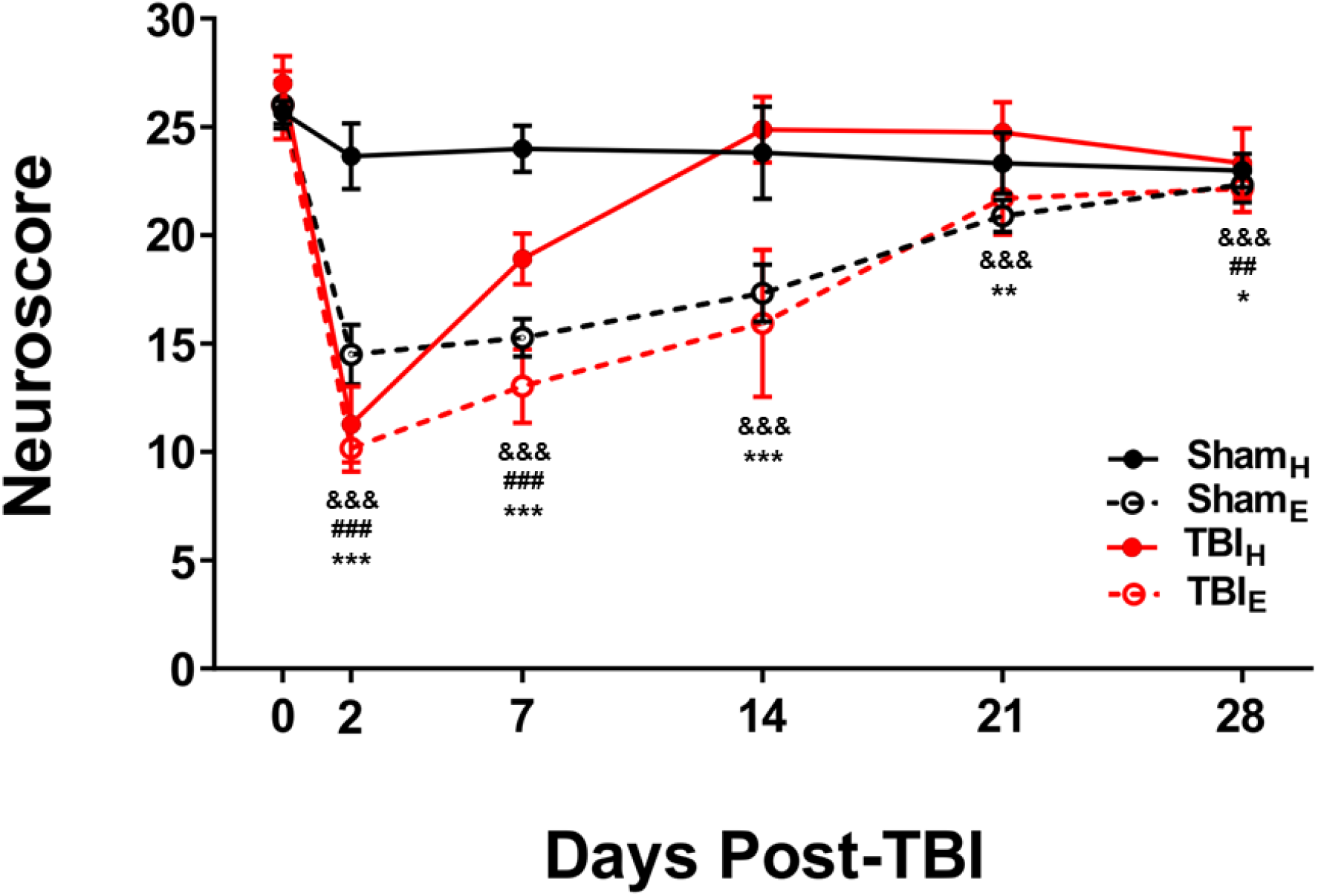
Neuroscore values shows the progression of sensorimotor deficits and recovery in each of the different groups. Notice the decreased neuroscore values 2 days after surgery in TBIE and TBIH, as well as ShamE rats, but not ShamH rats. TBIH rats neuroscore returned to baseline values faster than the TBIE and ShamE rats. Data are presented as mean ± SD. All the data were compared vs their baseline value (i.e. Day 0 score). ^&&&^p≤0.001 vs ShamE baseline value; ^##^p≤0.01, ^###^p≤0.001 vs TBIH baseline value; *p≤0.05, **p≤0.01, ***p≤0.001 vs TBIE baseline value, Bonferroni test corrected for multiple comparisons.

### Acute Electrographic posttraumatic seizures

EEG during the first 7 days after surgery recorded electrographic seizures, consisting of repetitive epileptiform discharges that evolved in frequency, amplitude, and morphology of discharges, and regularly ended with post-ictal amplitude suppression (Fig. 6). Two of six ShamH rats had a total of 4 seizures within 48 h after surgery and none thereafter. In these two rats, the mean latency to the first electrographic seizure was 19.5 ± 16.1 h, the mean seizure duration was 83.3 ± 35.5 s, and the total time spent having electrographic seizures was 2.8 ± 1.6 min (Table 2, Fig. 7).

**Fig. 6.**
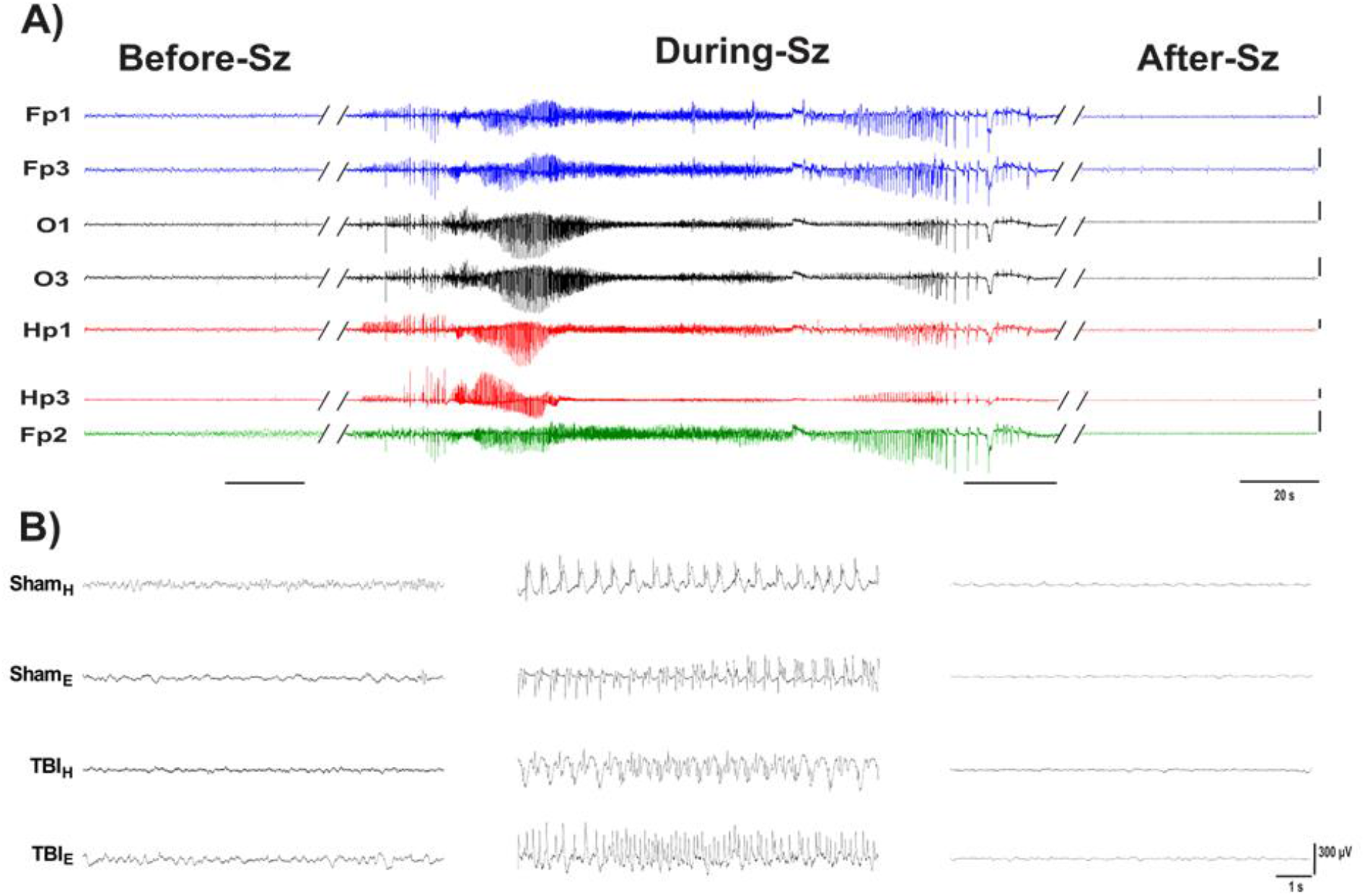
Acute electrograph seizures recorded from rats in each craniectomy group. (A): Example of acute generalized seizure recorded from one TBIH rat 2 days after surgery (EEG; 5-min epoch, high-frequency filter 90 Hz). Scale bar = 500 µV in each trace. (B) Expanded 10 s traces from a single channel (Fp1) taken 1 minute before the seizure onset (Before-Sz), at the mid-point of seizure (During-Sz) and 1 min after the seizure offset (After-Sz). Abbreviations: Sz; Seizure.

**Fig. 7.**
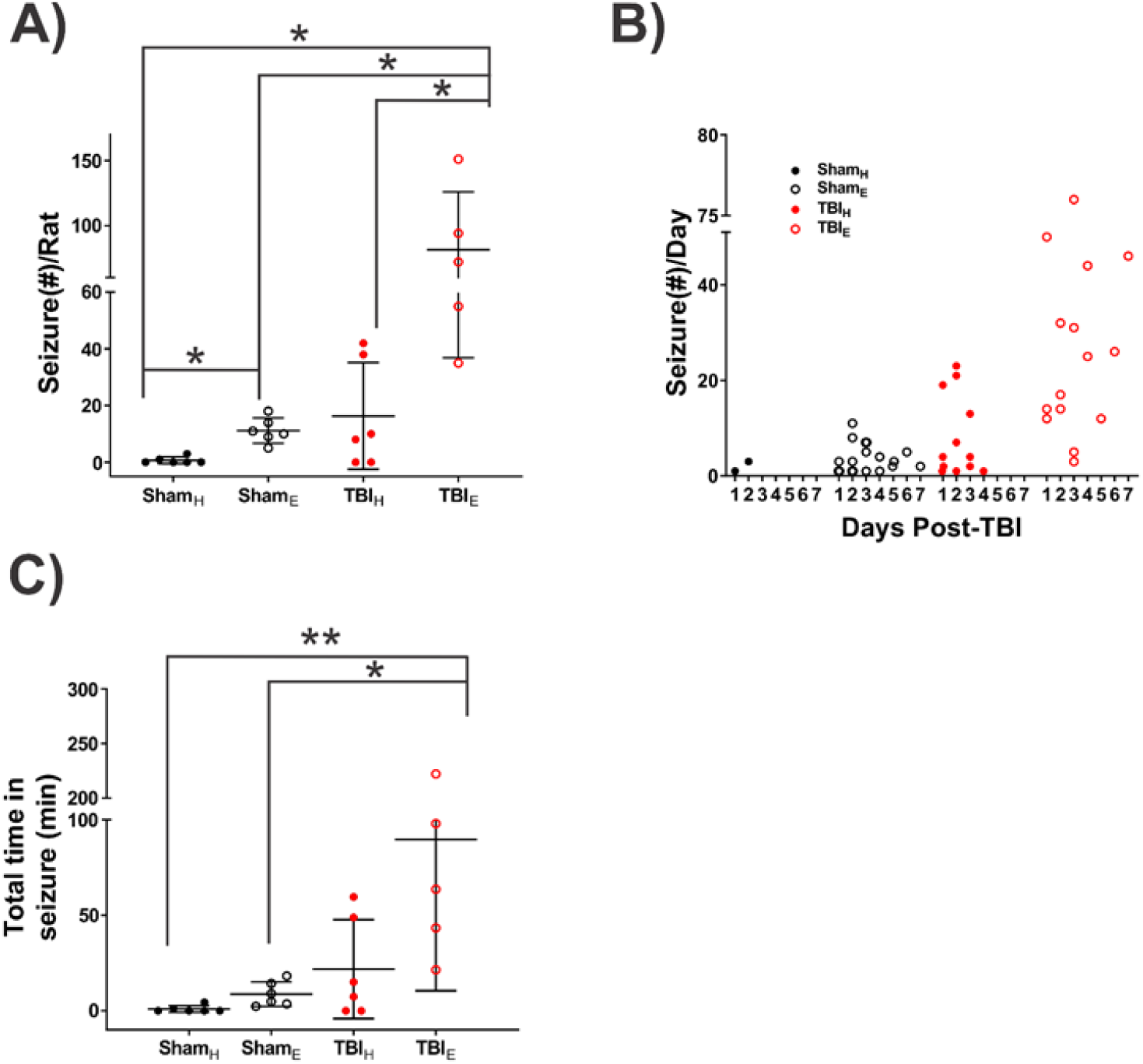
Quantification of acute EEG seizures during the first seven days after TBI A) Plot showing the number of seizures per rat after TBI. TBIE rats had significantly more acute EEG seizures than TBIH and ShamH rats. B) Number of EEG seizures per day for first 7 days. ShamE and TBIE rats had acute EEG seizures all 7 days, but ShamH and TBIH rat seizures resolved 2 and 4 days after surgery respectively C) TBIE rats spent significantly more time having seizures than ShamE and ShamH rats, but similar amount of time in seizures as TBIH rats. Data are presented as mean ± SD. Statistical significance: *p < 0.05; **p < 0.01; One-Way ANOVA test followed by post hoc analysis with Bonferrioni.

**Table 2.** Acute EEG seizures after TBI.

| **Group** | **EEG Seizure Activity** | **Latency (h)** | **Cumulative Seizure** | **Seizure Duration (s)** |
| --- | --- | --- | --- | --- |
| **Sham<sub>H</sub>** | 2/6 (33.3%) | 19.5±16.1 | 4 | 83.3±35.5 |
| **Sham<sub>E</sub>** | 6/6 (100%)* | 27.3±5.6 | 67 | 35.8±29.3 |
| **TBI<sub>H</sub>** | 4/6 (66.6%) | 12±1.6 | 98 | 79.9±56.5<sup>&&,#</sup> |
| **TBI<sub>E</sub>** | 6/6 (100%)* | 29.6±39.1 | 407 | 63.9±42.9<sup>&</sup> |

\*p≤0.5 vs Sham<sub>H</sub>, Chi-square test. <sup>&&</sup>p≤0.01, <sup>&</sup>p≤0.05 vs Sham<sub>E</sub>; <sup>#</sup>p≤0.05 vs TBI<sub>E</sub>; One Way Anova-Bonferrioni

All six ShamE rats (p<0.05, compared with ShamH rats) had seizures and overall, 67 seizures were recorded. The average number of seizures per ShamE rat was significantly higher than in ShamH rats (11.1 ± 1.8 vs. 2.0 seizures; p<0.05). Unlike the ShamH rats, ShamE rat seizures were detected during the entire first week of recording. The mean latency to the first seizure was 27.3 ± 5.6 h, the mean seizure duration was 35.8 ± 29.3 s, and the total time having electrographic seizures was 8.7 ± 2.6 min (Table 2, Fig. 7).

Four out of six TBIH rats had a total of 98 seizures during the first 96 h after LFPI and none in the remaining 72 h of the recording. The mean latency to the first seizure was shorter than latencies in ShamH and ShamE rats (12 ± 1.6 h), mean seizure duration was 79.9 ± 56.5 s, and total time having seizures was 32.7 ± 12.7 min (Table 2, Fig. 7).

All six TBIE rats had seizures (p<0.05, compared with ShamH group) that totaled 407 seizures during the first week of recording. There was a greater number of seizures per rat (81.4 ± 19.9 seizures), longer mean seizure duration (63.9 ± 42.9 s; p<0.05), and longer amount of time spent having electrographic seizures in TBIE rats than in ShamH and ShamE rats (89.7 ± 35.4 min; Table 2, Fig. 7).

## Discussion

The current study found greater MRI positive tissue deformation at 9 days and in most cases, weaker functional connectivity in the ipsilateral hemisphere up to 7 days after a craniectomy performed with an electrical drill than a hand drill in sham-injured rats. In TBI rats with an electric-drilled craniectomy, rats had prolonged sensorimotor deficits and greater number of acute seizures than TBI rats with hand-drilled craniectomy. Results suggest an electrical drill craniectomy can produce harmful brain structural and functional changes that worsens behavioral outcome and acute seizures after LFPI. Thus, the craniectomies conducted with an electric drill can constitute an injury itself which will doubtless reduce the effect size when comparing TBIs to sham controls.

Studies in rabbits found a craniectomy performed with electrical drill generates local heat that could be transmitted to the surface of the brain^26^. The heating and vibration from the electric drill could disrupt blood vessels and nerves that initiates an immediate primary neurovascular injury, followed by a more severe secondary biochemical cascade producing local cytotoxic edema^26,27^. The effects of heating could be reduced by continuous application of cold saline solution, regular breaks between drilling periods, and low drilling speeds combined with safest feed rates that minimizes the risks of penetrating the brain tissue^18^. However, it’s more difficult to dampen vibration. In contrast, the hand drill method generates the craniectomy by slow, steady, grinding pressure to penetrate the calvaria.

The mechanical stress induced by the electrical drill craniectomy likely contributes to the increased MRI positive tissue deformation after 9d. The tissue deformation could be related to edema, which is one of the primary lesions detected after injury and only partially resolves 7 days post-injury^28–31^. Whether edema is complication of intracerebral hemorrhage (ICH) and consequent increase in intracranial pressure is unclear^31^, though none of the animals showed signs of ICH (e.g., unresponsiveness to stimuli, immobility, lateral recumbent position, significant weight loss, etc.), but many had seizures. Earlier studies found T2 evaluation of the cortical lesion 3 days after the injury is a sensitive parameter to predict severity of histologically verified cortical neurodegeneration and functional impairment^24^. Thus, the acute MRI changes in animals with an electric drilled craniectomy could forecast long-term structural changes like gliosis, neuronal cell loss, synaptic reorganization. More recent reports showed the craniectomy procedure in sham animals induced the over expression of endothelial tyrosine kinase^32^, increase of KC-GRO and IFN-γ cytokines^27^ and imbalance in the pyruvate dehydrogenase kinase and pyruvate dehydrogenase phosphatase, which often correspond with alterations in the glucose metabolism and cellular energetics^33^. These latter changes could contribute to long-term morphological brain damage, impaired sensorimotor responses, and possibly increased brain excitability in sham animals^27^.

Severity of brain injury in animals with TBI also is assessed with sensorimotor tests like the neuroscore^34^. In the current study, we expected TBI rats to have lower neuroscore values that represent sensorimotor impairment 2d post-injury^35^. However, TBIH rats recovered faster than TBIE rats, and unexpectedly animals with electrical drilled craniectomy - TBI and sham - had the longest recovery and neuroscore values that were below baseline values 28 days after injury. Consistent with the MRI results, differences in neuroscores between animal groups suggest an electrical drilled craniectomy could produce more severe, widespread injury associated with functional deficits.

We performed an EEG analysis using GEC, which previous studies showed was a reliable and stable measure of EEG connectivity areas^36,37^, and recent work suggests the increased strength of GEC correlates with increased synchrony of inhibitory activity ^38^. In the current study, overall, there was weaker GEC in ShamE rats than ShamH rats. One interpretation of these results electrical drilling produces neuronal injury that includes a decrease in the synchrony of inhibitory activity ^39,40^. If this is correct, then a consequence of decreased inhibitory synchrony could be increased cortical excitability. Results from our analysis of early seizures support this interpretation since we detected the greatest number of seizures in TBIE rats and the number of rats with seizure and the number of seizures was comparable between ShamE and TBIH rats. The lowest number of seizures was in the ShamH rats. In this latter group, we detected seizures in two rats, and this could be due to the hand drilled craniectomy, though we cannot exclude the possibility that electrode implantation could induce acute injury and increased excitability ^41,42^. There is the implication of choice of craniectomy versus anesthetic control shams, which is especially relevant in studies of TBI. If extra-steps are not taken with electrical drilling, then an additional injury as described in the current study could confound studies of TBI, such as studies relating changes in MRI, electrophysiology, or behavior with severity of injury, which often is equated with force of impact or angle of pendulum. In our study, neuroscore appeared a good measure of total injury severity when using electric-drilled craniectomy. Also, choosing to control for the effect of craniectomy after LFPI by using shams with a craniectomy could affect biomarker studies of PTE. For example, an increase in acute seizures due to an injurious craniectomy could increase brain pathology and activity-dependent molecular changes that are related to craniectomy and not the TBI.

## Authorship contribution statement

**C. Santana-Gomez:** Methodology, Data curation, Formal Analysis, Visualization, Writing.

**G. Smith:** Methodology, Data curation. **A. Mousavi:** Data curation. **M. Shamas:** Data curation, Software. **N.G. Harris:** Project administration, Writing, Funding. **R. Staba:** Conceptualization, Writing, Project administration, Funding.

Author Disclosures: The authors have nothing to disclose.

## References

1. Thompson HJ, Lifshitz J, Marklund N, et al. Lateral fluid percussion brain injury: A 15-year review and evaluation. J Neurotrauma 2005;22(1):42–75; doi: 10.1089/neu.2005.22.42.

2. Xiong Y, Mahmood A, Chopp M. Animal models of traumatic brain injury. Nat Rev Neurosci 2013;14(2):128–142; doi: 10.1038/nrn3407.

3. Kabadi S V., D. HG, Stoica BA, et al. Fluid-percussion–induced traumatic brain injury model in rats. Nat Protoc 2010;5(9):1552–1563; doi: 10.1038/jid.2014.371.

4. Hayes RL, Stalhammar D, Povlishock JT, et al. A new model of concussive brain injury in the cat produced by extradural fluid volume loading: I. Biomechanical properties. Brain Inj 1987;1(1):73– 91.

5. Millen JE, Glauser FL, Fairman RP. A comparison of physiological responses to percussive brain trauma in dogs and sheep. J Neurosurg 1985;62(4):587–591; doi: 10.3171/jns.1985.62.4.0587.

6. Pfenninger EG, Reith A, Breitig D, et al. Early changes of intracranial pressure, perfusion pressure, and blood flow after acute head injury. Part 1: An experimental study of the underlying pathophysiology. J Neurosurg 1989;70(5):774–779; doi: 10.3171/jns.1989.70.5.0774.

7. Thibault LE, Meaney DF, Anderson BJ, et al. Biomechanical Aspects of a Fluid Percussion Model of Brain Injury. J Neurotrauma 1992;9(4):311–322; doi: 10.1089/neu.1992.9.311.

8. Härtl R, Medary MB, Ruge M, et al. Early white blood cell dynamics after traumatic brain injury: Effects on the cerebral microcirculation. Journal of Cerebral Blood Flow and Metabolism 1997;17(11):1210–1220; doi: 10.1097/00004647-199711000-00010.

9. McIntosh TK, Vink R, Noble L, et al. Traumatic brain injury in the rat: characterization of a midline fluid-percussion model. Neuroscience 1989;28(1):233–244; doi: 10.1016/0306-4522(89)90247-9.

10. Dixon CE, Lighthall JW, Anderson TE. Physilogic, histopathologic, and cineradiographic characterization of new fluid-percussion model of experimental brain injury in the rat. J Neurotrauma 1988;5(2):91–104.

11. Kharatishvili I, Pitkänen A. Association of the severity of cortical damage with the occurrence of spontaneous seizures and hyperexcitability in an animal model of posttraumatic epilepsy. Epilepsy Res 2010;90(1–2):47–59; doi: 10.1016/j.eplepsyres.2010.03.007.

12. Shultz SR, Cardamone L, Liu YR, et al. Can structural or functional changes following traumatic brain injury in the rat predict epileptic outcome? Epilepsia 2013;54(7):1240–1250; doi: 10.1111/epi.12223.

13. Ndode-Ekane XE, Santana-Gomez C, Casillas-Espinosa PM, et al. Harmonization of lateral fluid-percussion injury model production and post-injury monitoring in a preclinical multicenter biomarker discovery study on post-traumatic epileptogenesis. Epilepsy Res 2019;151(October 2018):7–16; doi: 10.1016/j.eplepsyres.2019.01.006.

14. Andrade P, Nissinen J, Pitkänen A. Generalized seizures after experimental traumatic brain injury occur at the transition from slow-wave to rapid eye movement sleep. J Neurotrauma 2017;34(7):1482–1487; doi: 10.1089/neu.2016.4675.

15. Kosaras B, Jakubowski M, Kainz V, et al. Sensory innervation of the calvarial bones of the mouse. Journal of Comparative Neurology 2009;515(3); doi: 10.1002/cne.22049.

16. Wu JCC, Chen KY, Yo YW, et al. Different sham procedures for rats in traumatic brain injury experiments induce corresponding increases in levels of trauma markers. Journal of Surgical Research 2013;179(1):138–144; doi: 10.1016/j.jss.2012.09.013.

17. Cole JT, Yarnell A, Kean WS, et al. Craniotomy: True sham for traumatic brain injury, or a sham of a sham? J Neurotrauma 2011;28(3):359–369; doi: 10.1089/neu.2010.1427.

18. Shoffstall AJ, Paiz JE, Miller DM, et al. Potential for thermal damage to the blood-brain barrier during craniotomy: implications for intracortical recording microelectrodes. J Neural Eng 2018;15(3); doi: 10.1088/1741-2552/AA9F32.

19. Olesen SP. Leakiness of rat brain microvessels to fluorescent probes following craniotomy. Acta Physiol Scand 1987;130(1):63–68; doi: 10.1111/j.1748-1716.1987.tb08112.x.

20. Hatashita S, Koike J, Sonokawa T, et al. Cerebral edema associated with craniectomy and arterial hypertension. Stroke 1985;16(4):661–668; doi: 10.1161/01.STR.16.4.661.

21. Paxinos G, Watson C. The Rat Brain in Stereotaxic Coordinates. 6th edn. Academic Press/Elsevier: Amsterdam ; Boston; 2007.

22. Tustison NJ, Avants BB, Cook PA, et al. N4ITK: improved N3 bias correction. IEEE Trans Med Imaging 2010;29(6):1310–1320; doi: 10.1109/TMI.2010.2046908.

23. Pitkanen A, McIntosh T. Animal models of post-traumatic epilepsy. Journal of Neurot 2006;23(2):241–261.

24. Kharatishvili I, Sierra A, Immonen RJ, et al. Quantitative T2 mapping as a potential marker for the initial assessment of the severity of damage after traumatic brain injury in rat. Exp Neurol 2009;217(1):154–164; doi: 10.1016/j.expneurol.2009.01.026.

25. Gotman J. Automatic seizure detection: improvements and evaluation. Electroencephalogr Clin Neurophysiol 1990;76(4):317–324; doi: 10.1016/0013-4694(90)90032-F.

26. Edvinsson L, West KA. Experimental Cerebral Heat Lesions Produced By Trephine Craniotomy in Rabbits. Acta Pathologica Microbiologica Scandinavica Section A Pathology 1972;80 A(1):134–138; doi: 10.1111/j.1699-0463.1972.tb00278.x.

27. Cole JT, Yarnell A, Kean WS, et al. Craniotomy: True sham for traumatic brain injury, or a sham of a sham? J Neurotrauma 2011;28(3):359–369; doi: 10.1089/neu.2010.1427.

28. Soares HD, Thomas M, Cloherty K, et al. Development of Prolonged Focal Cerebral Edema and Regional Cation Changes Following Experimental Brain Injury in the Rat. J Neurochem 1992;58(5):1845–1852; doi: 10.1111/j.1471-4159.1992.tb10061.x.

29. Onyszchuk G, He YY, Berman NEJ, et al. Detrimental effects of aging on outcome from traumatic brain injury: A behavioral, magnetic resonance imaging, and histological study in mice. J Neurotrauma 2008;25(2):153–171; doi: 10.1089/neu.2007.0430.

30. Szczygielski J, Glameanu C, Müller A, et al. Changes in posttraumatic brain edema in craniectomy- selective brain hypothermia model are associated with modulation of aquaporin-4 level. Front Neurol 2018;9(OCT):1–16; doi: 10.3389/fneur.2018.00799.

31. Assaf Y, Beit-Yannai E, Shohami E, et al. Diffusion- and T2-weighted MRI of closed-head injury in rats: A time course study and correlation with histology. Magn Reson Imaging 1997;15(1):77–85; doi: 10.1016/S0730-725X(96)00246-9.

32. Wu JCC, Chen KY, Yo YW, et al. Different sham procedures for rats in traumatic brain injury experiments induce corresponding increases in levels of trauma markers. Journal of Surgical Research 2013;179(1):138–144; doi: 10.1016/j.jss.2012.09.013.

33. Xing G, Ren M, O’Neill JT, et al. Controlled cortical impact injury and craniotomy result in divergent alterations of pyruvate metabolizing enzymes in rat brain. Exp Neurol 2012;234(1):31– 38; doi: 10.1016/j.expneurol.2011.12.007.

34. Niskanen JP, Airaksinen AM, Sierra A, et al. Monitoring functional impairment and recovery after traumatic brain injury in rats by fMRI. J Neurotrauma 2013;30(7):546–556; doi: 10.1089/neu.2012.2416.

35. Ndode-Ekane XE, Santana-Gomez C, Casillas-Espinosa PM, et al. Harmonization of lateral fluid-percussion injury model production and post-injury monitoring in a preclinical multicenter biomarker discovery study on post-traumatic epileptogenesis. Epilepsy Res 2019;151; doi: 10.1016/j.eplepsyres.2019.01.006.

36. Bragin A, Almajano J, Kheiri F, et al. Functional connectivity in the brain estimated by analysis of gamma events. PLoS One 2014;9(1):1–8; doi: 10.1371/journal.pone.0085900.

37. Kheiri F, Bragin A, Engel JJ. Functional connectivity between brain areas estimated by analysis of gamma waves _ Elsevier Enhanced Reader.pdf. J Neurosci Methods 2013;214:184–191.

38. Shamas M, Yeh HJ, Fried I, et al. Disorders of the Nervous System Interictal Gamma Event Connectivity Differentiates the Seizure Network and Outcome in Patients after Temporal Lobe Epilepsy Surgery. 2022; doi: 10.1523/ENEURO.0141-22.2022.

39. Rachmany L, Tweedie D, Rubovitch V, et al. Cognitive impairments accompanying rodent mild traumatic brain injury involve p53-dependent neuronal cell death and are ameliorated by the tetrahydrobenzothiazole PFT-α. PLoS One 2013;8(11):1–12; doi: 10.1371/journal.pone.0079837.

40. Benady A, Freidin D, Pick CG, et al. GM1 ganglioside prevents axonal regeneration inhibition and cognitive deficits in a mouse model of traumatic brain injury. Sci Rep 2018;8(1):1–10; doi: 10.1038/s41598-018-31623-y.

41. Andrade P, Banuelos-Cabrera I, Lapinlampi N, et al. Acute Non-Convulsive Status Epilepticus after Experimental Traumatic Brain Injury in Rats. J Neurotrauma 2019;36(11):1890–1907; doi: 10.1089/neu.2018.6107.

42. Fox R, Santana-Gomez C, Shamas M, et al. Title: Different Trajectories of Functional Connectivity Captured with Gamma-Event Coupling and Broadband Measures of EEG in the Rat Fluid Percussion Injury Model. Running Title: EEG Functional Connectivity after Rat FPI. n.d.; doi: 10.1101/2024.06.02.597056.

